# Neuroinfectiology of an atypical anthrax-causing pathogen in wild chimpanzees

**DOI:** 10.1101/2023.11.21.568053

**Authors:** Tobias Gräßle, Carsten Jäger, Evgeniya Kirilina, Jenny E. Jaffe, Penelope Carlier, Andrea Pizarro, Anna Jauch, Katja Reimann, Ilona Lipp, EBC consortium, Roman M. Wittig, Catherine Crockford, Nikolaus Weiskopf, Fabian H. Leendertz, Markus Morawski

## Abstract

*Bacillus cereus* biovar *anthracis* (*Bcbva*) is an atypical anthrax-causing bacterium, inflicting wildlife fatalities across African rainforest ecosystems. The pathogen’s virulence in one of our closest living relatives, the chimpanzee, together with human serological evidence, suggests *Bcbva* is zoonotic. While classical *B. anthracis*-induced anthrax has been described to affect the central nervous system at a progressive disease-state, the neuroinfectiology of *Bcbva* is yet unknown. Here we characterised the pathogen’s neuro-invasiveness via gross pathological assessment, ultra-high resolution quantitative Magnetic Resonance Imaging and histological analysis on four brains, which were extracted from naturally deceased wild chimpanzees in Taï National Park, Côte d’Ivoire.

Based on macroscopically evident pial vessel congestion and haemorrhages as well as cortical siderosis detected via MRI, we concluded that *Bcbva* induced meningitis analogous to *B. anthracis*. Further, histological visualisation of bacteria and leukocytes in the subarachnoid space evidenced the bacterium’s capability to breach the arachnoid barrier. *Bcbva* was detected in the brain parenchyma of all four cases. This indicates a higher ability to transgress the glia limitans and therefore exhibits a higher neuroinvasiveness compared to *B. anthracis* that predominantly stays confined to the meninges. Heightened glial fibrillary acidic protein (GFAP) expression but little morphological gliosis suggest a rapid disease progression leading to host-death within hours to a few days after central nervous system invasion.

Overall our results reveal *Bcbva*’s ability to breach blood-brain barriers which results in a pronounced neuropathogenicity. *Bcbva* causes extensive damage to the meninges and the brain parenchyma, as well as rapid and massive digestion of brain extracellular matrix in chimpanzees and potentially so in humans in case of zoonotic spillover.

## Introduction

The recent COVID-19 pandemic has showcased the significant socio-economic costs caused by neuropathogenic effects of zoonotic diseases^1,2^. Understanding the neuropathological mechanisms of these diseases before outbreaks is crucial for the development of therapies and diagnostics^3^. This presents a challenge, as common laboratory animals do not always serve as ideal models for the complexity of the large human brain. Hierin, we demonstrate that unique insights into the neuroinfectiology of a likely zoonotic disease can be gained by studying naturally deceased wild chimpanzees in an opportunistic, ethical approach.

It was unexpected when in the early 2000s deaths of wild chimpanzees in the dense west African rainforest of Taï National Park were attributed to anthrax^4^, as this lethal, bacterial, zoonotic disease usually affects ungulates in savannahs or other open landscapes^5^. Genetic investigations revealed that the bacterium found in these chimpanzees was phylogenetically distinct to the monophyletic *B. anthracis* lineage, which thus far had been believed to be the only agent capable of inducing anthrax^6^. The newly discovered pathogen was more closely related to *Bacillus cereus* strains – bacteria usually associated with food poisoning^7^. New anthrax-inducing strain was termed *Bacillus cereus* biovar *anthracis* (*Bcbva*)^8^. Since its discovery, *Bcbva* has been associated with wildlife fatalities across west and central African rainforests, distinct from the ecological niche and geographical distribution of *B. anthracis*^9,10^. It predominantly affects primates, including apes and old world monkeys, followed by forest antelopes and accounts for up to 40% of the observed wildlife mortality locally^10^.

On a molecular level, *B. anthracis* and *Bcbva* share highly syntenic virulence plasmids, decisive for both bacteria’s pathogenicity^11^. The larger of these plasmids, pXO1 (182 kilobase pairs (kbp)) encodes for the anthrax toxin complex responsible for disrupting the host’s innate immune response, while the smaller plasmid, pXO2 (96 kbp) encodes a poly-γ-D-glutamic acid capsule operon, shielding vegetative bacteria from phagocytes and antibody binding^12,13^.

Although no confirmed human *Bcbva*-case has been reported yet, *Bcbva*’s capability to elicit lethal infections in our closest living relatives, paired with human serological evidence, strongly vouch for *Bcbva*’s zoonotic character^14^ .

*B. anthracis* morbidity in humans is predominantly of zoonotic origin, induced by contact with spore-contaminated animal products. Therefore, bushmeat poses a serious potential source of human *Bcbva* infection^15^. In addition to zoonotic infections, *B. anthracis* has become notorious for its misuse in biological warfare and bioterrorism^16,17^.

*B. anthracis* is known to be neuroinvasive and meningitis is a common severe complication of systemic anthrax^18,19^. Involvement of the central nervous system (CNS) poses a clear negative prognostic factor for human survival and mortality rates are close to 100 % in spite of aggressive medical therapy^20,21^. The brief, rapidly progressing course of disease is characterised by nausea, emesis, severe headache, meningeal signs, seizures and other neurological deficits shortly prior to death^22,23^. The cerebrospinal fluid (CSF) regularly presents haemorrhagic with a marked leukocytosis and bacterial infiltration. MRI findings include leptomeningeal enhancement, vasogenic oedema^24^, abscessation^25^, multiple infarction and foremost haemorrhages^26^. In general, the inflammatory pattern in the CNS analogous to other affected organs, is of a predominantly haemorrhagic character. In severe cases, a diffusely crimson coloured brain surface is revealed upon *postmortem* examination which has been metaphorically coined “cardinal’s cap”^27^.

Thus far, nothing was known about the neurovirulence of *Bcbva* in its natural hosts and particularly in humans. We addressed this knowledge gap by analysing four brains extracted from wild chimpanzees that died of *Bcbva*-induced anthrax in Taï National Park. Chimpanzees are one of humans’ closest living relatives providing the best model to study *Bcbva*’s neuroinfectology in the case of zoonotic spillover. In the late 1950s, chimpanzees were used as a primate animal model for inhalational anthrax infection in experiments with intended multiple aerosol infections^28^. Fortunately, the progress in animal welfare in laboratory research has recently restricted the use of great apes in invasive medical research^29^, which is banned by an ever growing number of scientific organisations worldwide^30^. As brain specimens in our study were obtained occasionally after the chimpanzees’ natural deaths, providing an ethical approach to study the pathogen’s neurovirulence^31^.

The analyses reveal the bacterium’s pronounced neuroinvasiveness, enabling it to disrupt blood-brain barriers with subsequent propagation in the meninges and brain parenchyma as well as severe effects on brain tissue integrity.

## Materials and methods

### Cases

Four wild chimpanzee (*Pan troglodytes verus*) brains, with confirmed fatal *Bcbva* infections, from Taï National Park (Taï NP, Côte d’Ivoire) were examined, along with one control specimen from Tacugama Chimpanzee Sanctuary (Sierra Leone), which died after an intra-group attack. Details on age, sex, and *postmortem* interval before fixation (PMI) are displayed in Supplementary Table 1 and summarised below. Our research is conducted according to standards specified by the Max Planck Society’s Ethics Committee (04.08.2014). Our *postmortem* brain extraction methods are noninvasive in the sense that they do not harm the living animal.

### Behaviour

Chimpanzees in Taï NP are the subjects of behavioural research conducted within the <u>Taï</u> <u>Chimpanzee Project</u> (TCP)^32^. Habituated chimpanzees in Taï NP are followed daily by a research team, and their behaviour and health status are systematically documented within ongoing health monitoring with continuous presence of a field veterinarian as an integral part of TCP.

### Necropsy

*Postmortem* examinations were a part of the ongoing wildlife health monitoring programme established in 2001^32^ aiming to understand and monitor the drivers of wildlife mortality, especially zoonotic diseases relevant for species conservation and public health^33^.

*Postmortem* tissue sampling was performed exclusively by trained veterinarians following strict biosafety protocols^31^. The carcasses were solely approached wearing protective equipment consisting of rubber boots, a Tyvek® coverall, three pairs of gloves, protective arm sleeves, an FFP3 mask and a face shield/goggles. All single-use equipment utilised during the necropsy was enclosed in at least two layers of trash bags and was subsequently burnt. Reusable material and sample containers were decontaminated at a hygiene barrier using bleach with a sodium hypochlorite concentration of 1%. Brain extractions were performed utilising a battery-operated oscillating saw (KUGEL medical) and the complete organ was immersed in 10% neutral buffered formalin. Tissue samples for molecular biology diagnostics were placed into cryo tubes (Simport) and stored in liquid nitrogen. After export to Germany, the brains were transferred into a phosphate-buffered saline (PBS) pH 7.4 solution with 0.1% sodium azide for long-term storage at 4°C. For the complete detailed sampling protocol please refer to^31^. The meninges were removed prior to the start of the consequent investigations due to requirements of a different project.

### Genetics

#### PCR Diagnostics

DNA for PCR-based diagnostics was extracted from tissue samples with the Viral RNA Mini Kit (Qiagen©), following the manufacturer’s recommended protocol. The PCR master mix was prepared utilising Platinum™Taq DNA Polymerase and DNA-free reagents (Invitrogen), along with primer-pairs targeting three *Bcbva* markers (one for each virulence plasmid and one chromosomal) and a corresponding, fluorophore-labelled TaqMan probe for real-time detection (Supplementary Table 2)^10^. Five µl or a maximum of 200 ng/µl of template DNA was added per well, and the assay was run on an AriaMx Real-time PCR System (Agilent). The cycle conditions were as follows: 95 °C for 10 min, followed by 45 cycles of 95 °C for 15 s and 60 °C for 34 s. Each sample was tested in duplicate, and duplicate negative template controls and standard dilutions were included. The amplification data were analysed with the Agilent AriaMx v. 1.7 software.

#### Bacterial culture

Spleen tissue samples were cultured on R&F® *Bacillus cereus*/*Bacillus thuringiensis* Chromogenic Plating Medium (R & F Products©) via dilution streaking and single colonies were picked after 14 h of incubation. Bacterial DNA was extracted utilising the DNeasy Blood & Tissue Kit (QIAGEN©) with the manufacturer’s recommended protocol for gram positive bacteria. The DNA concentrations of extracts were quantified with the help of Qubit™ dsDNA HS Assay Kit (Invitrogen) and 1 ng of genomic DNA was used for constructing sequencing libraries with the NexteraXT Kit (Illumina). MagSi-NGS^PREP^ Plus Beads (Steinbrenner Laborsysteme GmbH) were employed for PCR clean-up with a volume ratio of 0.7 (beads/amplified library) and freshly prepared 80% ethanol. Libraries were sequenced on a NextSeq 2000 platform using the P2 reagents (Illumina).

Previously published *Bcbva* whole-genome sequence data^10^ were downloaded from the sequence read archive (SRA) and converted to the fastq format with the SRA toolkit v. 2.22.0^34^.

#### Bioinformatics and phylogenetic analysis

Basecalling and demultiplexing of the newly sequenced chimpanzee isolates was performed via bcl2fastq v. 2.19.0.316. Resulting fastq data were quality assessed and adapter trimmed with fastp v.0.20.1^35^. Genetic variants were called using the snippy pipeline^36^ with default parameters and variants were annotated by means of SnpEff v.5.1^37^. A SNP-alignment was extracted via snp-sites v. 2.5.1^38^ and a suitable DNA-evolution model for phylogenetic inference was chosen utilising modeltest-ng v.0.1.7^39^. The chromosomal *Bcbva* phylogenetic tree was built in a maximum likelihood framework using RAxML-NG v.1.1^40^, starting from 20 random and 20 parsimony trees. The best scoring ML tree was resampled with 500 bootstrap replicates to calculate transfer bootstrap expectancy (TBE) branch support values^41^. The resulting tree was imported into RStudio^42^ via the treeio package^43^ and visualised via ggtree^44^.

#### Ultra-high resolution quantitative MRI

Macroscopic neuropathological changes caused by *Bcbva* we accessed used ultra-high resolution quantitative MRI (qMRI), providing microstructural information about tissue pathology^45–47^. We obtained maps of three qMR parameters: magnetisation transfer saturation (MTsat) reflecting tissue macromolecular content, longitudinal relaxation rate (R1) reflecting myelin and iron content, effective transverse relaxation rate R2* strongly sensitive to tissue iron^46^.

QMRI imaging was performed on a human whole-body 7T MRI scanner (Terra, Siemens Healthineers, Erlangen), using a 32 channel radio-frequency (RF) transmit-receive human head coil (Nova Medical, Wilmington, MA). For MRI investigations the brains were embedded in the MRI-invisible liquid Fomblin (Solvay). QMRI was performed using a multi-parametric mapping method^45,47,48^. Scanning parameters are provided in^45^ and in Supplementary Materials. The quantitative maps of R1, MTsat and R2* with isotropic resolution of 300 µm were obtained using the open source hMRI toolbox^49^, adapted for the analysis of ultra-high resolution *postmortem* images^45^.

### Histology

The brains were cut into 15 mm thick coronal slabs and cryoprotected in 30% sucrose in PBS pH 7.4 with 0.1% sodium azide. Sections of 30 µm thickness were cut on a cryomicrotome (Microm HM430 with freezing unit Microm KS34, Thermo Scientific) and mounted on coated glass slides, air dried and rehydrated before further processing. Consecutive sections were stained using the haematoxylin and eosin (H/E) and Nissl stain^50,51^ for bacteria visualisation, Perls’s stain with diaminobenzidine enhancement for iron visualisation, and a modified Moeller, Rakette stain and the classical Gram-stains^50^ for detection of endospores.

All immunohistochemical stains were performed on free floating sections, which were treated with 2% H_2_O_2_ in 60% methanol for 30 min to reduce endogene peroxidase activity, followed by a blocking step with a solution containing 2% bovine serum albumin and 0.5% donkey serum. Sections were incubated for 24 or 48 h at 4°C with the primary antibodies (see Supplementary Table 3 for the list of all used materials). After incubation with secondary antibodies for 1 h at room temperature and for 1 h with ExtrAvidin-Peroxidase, the immunoreaction was detected through a diaminobenzidine reaction (partially with nickel enhancement) under visual control. Sections were counterstained with the Nissl stain. The sections were coverslipped with Entellan (Merck) and imaged with Zeiss AxioScan.Z1 (Zeiss, Jena, Germany).

#### Histological quantification

The degree of *Bcbva* invasion in Nissl-stained sections was assessed semi-quantitatively by one rater (CJ). The bacterial load in the section of each case was rated on the five level scale according to categories: “very sparse”, “sparse”, “moderate”, “dense”, “very dense”.

To assess *Bcbva*’s impact on parenchymal tissue integrity, perineuronal nets (PNNs) were stained via anti-aggrecan immunohistochemistry. PNNs are a specialised form of neuronal extracellular matrix (ECM) showing a mesh-like arrangement of proteins and sugar molecules ensheathing certain neurons, with aggrecan being a key component of healthy neurons^52^. The number of PNN-ensheathed neurons was quantified by counting the number of anti-aggrecan antibody-labelled neurons per mm^2^ in all cases (Table 1) by one rater (KR), blinded to the experimental conditions. The same cortical area (primary visual cortex, analogue of Brodmann Area 17) was analysed in all cases to account for known variation of density and cortical distribution of PNNs between cortical areas^51^. Counts were performed on a Zeiss AxioScan.Z1 (Zeiss, Jena, Germany) using Zen Blue software 3.6 (Zeiss, Jena, Germany). Relevant cortical regions were delineated at low magnification and three frames were placed in a systematic-consecutive fashion in the delineated regions. Neurons within each frame (250 x 2500 µm) were counted at higher magnification (20x, NA 0.8, ∼350nm). Three sections and six frames per case were investigated. The number of PNNs was converted to numerical density per mm^2^.

**Table 1:** Assessment of bacterial load and perineuronal nets’ integrity in different brain areas. The semiquantitative assessment was performed for iv=intravascular and ev=extravascular compartments on the following scale:; + very sparse; ++ sparse; +++ moderate; ++++ dense ; +++++ very dense and was validated by quantitative analysis on the occipital cortex. Numerical density of neurons with perineuronal nets decreased with the increase of *Bcbva* bacterial load in the primary visual cortex. All infected individuals showed a significantly lower PNN density compared to the control case, as demonstrated by an unpaired Student’s t-test, with p<.001 after family-wise-error correction.

|  | frontal | central | cerebellum | brainstem | occipital | quantitative validation<br>occipital cortex, V1<br>total *Bcbva* load<br>*Bcbva* bacteria per<br>1mm<sup>3</sup> brain tissue | # PNNs / mm<sup>2</sup> | *Bcbva*<br>covered<br>area / total<br>area [%] |
| --- | --- | --- | --- | --- | --- | --- | --- | --- |
| case1 | iv +++++<br>ev +++++ | iv ++++<br>ev ++++ | iv ++++<br>ev ++++ | iv +++++<br>ev ++++ | iv ++++<br>ev +++ | $2.2 \pm 1.1 \times 10^4$ | $1.01 \pm 1$ (*$p=0.5 \times 10^{-6}$) | 0.4 |
| case 2 | iv ++++<br>ev ++ | iv +++<br>ev ++ | no data | iv +++<br>ev ++ | iv +++<br>ev + | $0.6 \pm 0.3 \times 10^4$ | $42.4 \pm 4$ (*$p=0.1 \times 10^{-6}$) | 0.1 |
| case 3 | iv +++<br>ev + | iv +++<br>ev + | no data | iv +++<br>ev + | iv ++<br>ev + | $0.1 \pm 0.05 \times 10^4$ | $65 \pm 4$ (*$p=0.2 \times 10^{-5}$) | 0.02 |
| case 4 | no data | no data | no data | no data | iv ++<br>ev + | $0.05 \pm 0.25 \times 10^4$ | $85 \pm 11$ (*$p=0.001$) | 0.01 |
| control | - | - | - | - | - | 0 | $117 \pm 7.3$ | 0 |

*Bcbva* load was quantified on consecutive Nissl sections. Percentual area coverage with *Bcbva* was estimated using AI software module Intellesis Zen Blue 3.6. The algorithm was trained on manually labelled bacteria on ten randomly selected frames from all cases. The trained algorithm was applied to one section and six frames per case. The area (µm^2^) covered with *Bcbva* was converted to percentual *Bcbva* coverage in relation to total frame area.

We compared PNN density between each infected case and healthy control using unpaired two-sided t-tests for samples with different variance, comparing six frames of *Bcbva*-cases with the six samples from control brain, correcting for multiple comparisons (four comparisons corresponding to four cases) using the family-wise-error correction method.

The area measures were used to estimate the total density of *Bcbva* in bacteria per mm³ of brain tissue, by assuming non-overlapping bacteria and estimating the area covered by one bacteria to be about 6 µm^2^, based on the average bacterial length and width of 1 µm and 6 µm respectively.

### Data availability

High-throughput sequencing reads of the two novel *Bcbva* genomes are available via the short read archive. All additional data used in this manuscript may be shared upon request.

## Results

### Gross pathology indicates inflammatory meningeal involvement

Included in this study were four complete chimpanzee brains, sampled between December 2018 and March 2020 (Supplementary Table 1). Tissue samples from these individuals were incidentally found to be PCR-positive for all three *Bcbva* markers.

The necropsy on the young adult female case 1 (14±3 y) from an unhabituated group was conducted on 12th of December 2018 and, since it was an unknown individual, no prior behavioural history was available. The chimpanzee was assessed to have died less than 24 h prior to the necropsy as assessed via fly larvae infestation and *rigor mortis* status. The body was bloated and hyperaemic. Haemorrhagic lungs were the major findings on gross pathology.

Case 2 (1.7 y), a male infant, died suddenly on 5th of April 2019 without prior clinical signs up to the last observation 1.5 h before death and the brain was extracted 24 h after death. The main macroscopic findings included severe, haemorrhagic pneumonia and a congested liver, as well as a mild haemorrhagic meningitis with congested meningeal vessels, multifocal meningeal haemorrhages (up to 10 mm in diameter, see inset of Fig. 1 b for an example) and oedematous brain tissue.

**Fig.1.**
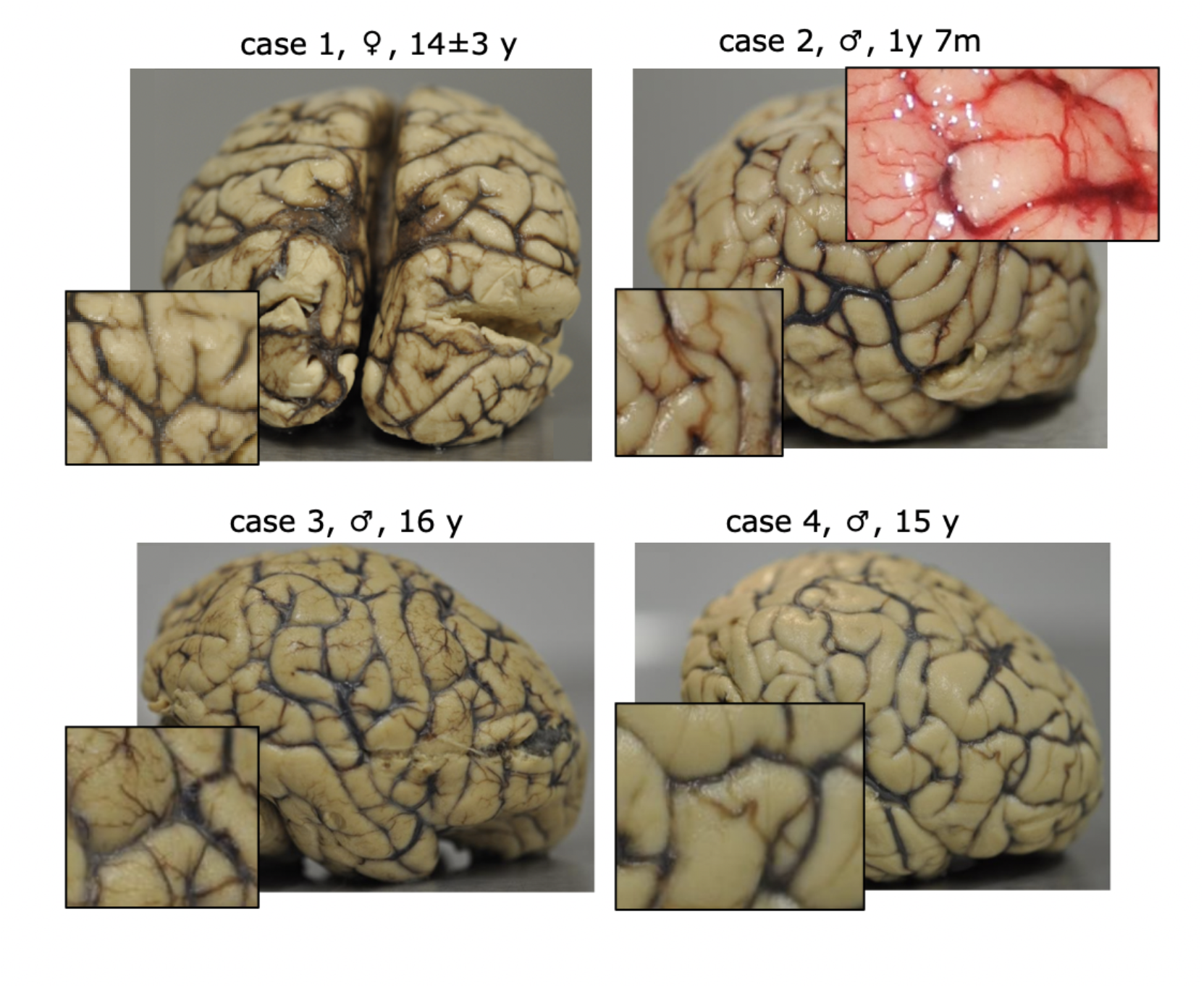
Postmortem formalin-fixed brains of all four *Bcbva* cases. The superficial vessels were congested to different degrees (most apparent in cases 1-3) pointing towards inflammatory meningeal involvement (inserts). For case 2, the photograph taken during brain extraction before fixation also shows focal meningeal haemorrhage (insert).

The two remaining specimens stem from case 3 (16 y) and case 4 (15 y), two socially bonded adult males. Case 3 was found freshly dead near his nesting site on the morning of 11th of March 2020 and the brain was extracted within 16 h after death. He was previously in good general condition without exhibiting any signs of disease when last observed five days prior to death. Case 4 interacted with case 3’s corpse on 11th of March 2020, repeatedly biting it and eventually creating a large, deep lesion in the throat area (∼25 x 20 cm). On 12th and 13th of March, he was more subdued, producing no panthoots or drums, and on the 13th he was resting much more than normal. On the 14th of March he separated from the group in the morning and was resting on his back approximately 70% of the time; his breathing and eating behaviour was evaluated as normal. He was found dead under his nesting-site on the morning of 15th of March and the brain was extracted within 12 h after death. Both cases 3 and 4 exhibited a severe haemorrhagic pneumonia on *post mortem* examination.

No gross brain pathology was reported in the necropsy records of case 1,3, and 4. However, based on photographs taken during brain fixation, the brains of cases 1 and 3 exhibited clearly injected and congested pial vessels, which corresponds to the gross-pathological findings of case 2 (see Fig. 1). Further, haemorrhages in the caudal part of the frontal lobe bilateral to the longitudinal fissure (10 x 10 mm) were still apparent in the meninges of case 1. These findings indicate inflammatory meningeal involvement to varying degrees.

### Whole-genome phylogenies and SNP-analysis reveal genetically representative *Bcbva* strains

*Bcbva* was successfully cultured from tissue samples of cases 3 and 4 and whole-genome sequences exceeding a 200X depth of coverage were generated. The isolates exhibited identical SNP profiles, indicating that the chimpanzee deaths were epidemiologically linked. It is possible that both chimpanzees were exposed to a common infection source, or that case 4 became infected while interacting with case 3’s carcass. Compared to the *Bcbva* reference genome (GCF_000143605.1), which likewise stems from Taï NP, the isolates exhibited 69 SNPs and fell well within the known diversity of Taï NP isolates (Fig. 2). Sixty-eight of those SNPs were chromosomal, whereas one was identified on the virulence plasmid pXO2. Among the 48 SNPs located in protein coding sequences, 38 were non-synonymous, resulting in four occasions in an early stop codon (SNP Supplementary Table 2). However, none of the SNPs were located in the virulence-defining pathogenicity islands of pXO1 and pXO2. Thus the isolates’ phylogenetic placements and SNP-profiles provide no evidence of altered (neuro)pathogenicity within these strains, indicating that they are representative for *Bcbva*’s neuroinvasive capabilities.

**Fig. 2.**
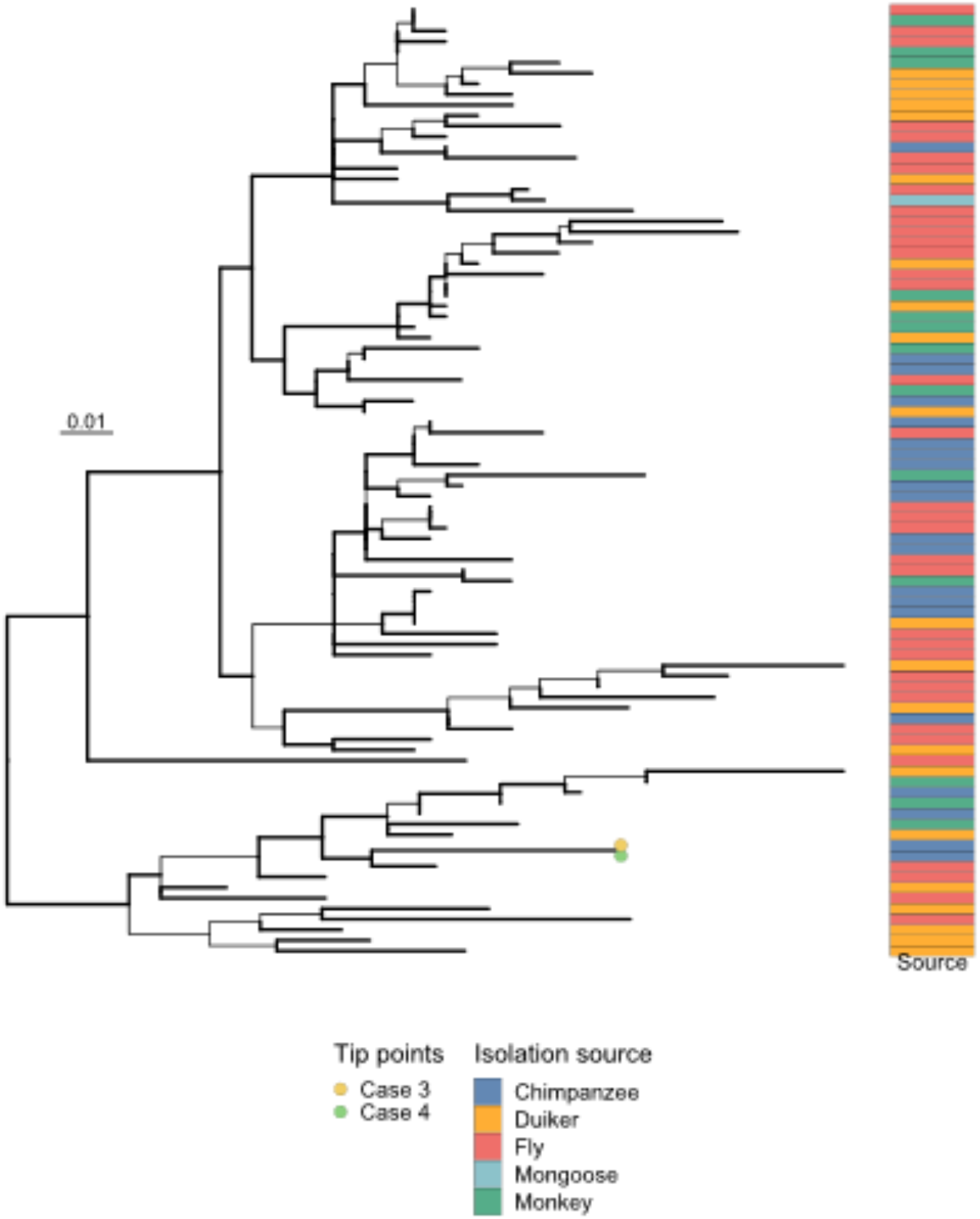
Chromosomal maximum likelihood SNP phylogeny of *Bacillus cereus* biovar *anthracis* (n = 90). The tree is midpoint rooted and internal branches with transfer bootstrap values below 0.7 are plotted in non-bold. Tip points indicate the isolates of case 3 and case 4, which fall within the diversity of previously published isolates. The isolation sources of all sequences are illustrated by a coloured strip and the scale bar indicates the substitutions per site. No evidence of altered (neuro)pathogenicity within the isolated strains was found, suggesting that the study cases are representative for *Bcbva’*s neuroinvasive capabilities.

### *Bcbva* induces superficial cortical siderosis and lytic lesions as revealed by qMRI

Maps of qMRI parameters R2*, R1 and MTsat for all cases with *Bcbva* infection demonstrated clear differences compared to the control (Fig. 3). Infected brains exhibited lytic lesions in white and grey matter. These lytic lesions, visible in all MRI contrasts, were probably the result of *Bcbva*-induced heterolysis. In three cases (case 1, 3 and 4), lytic processes additionally resulted in inhomogeneity of quantitative R1 maps with a hyperintense brain periphery and a hypointense brain centre. Therefore, the observed gradient of R1 likely reflects the disintegration of brain macromolecules such as lipids, proteins and ECM, caused by inhomogeneous fixation or by heterolysis^53,54^. Similar gradients of tissue integrity are often observed in immersion-fixed large brains (i.e. human brains)^55^. The fact that the gradient was more pronounced for *Bcbva*-infected brains as compared to the control may indicate that infection contributes to enhanced heterolysis.

**Fig.3.**
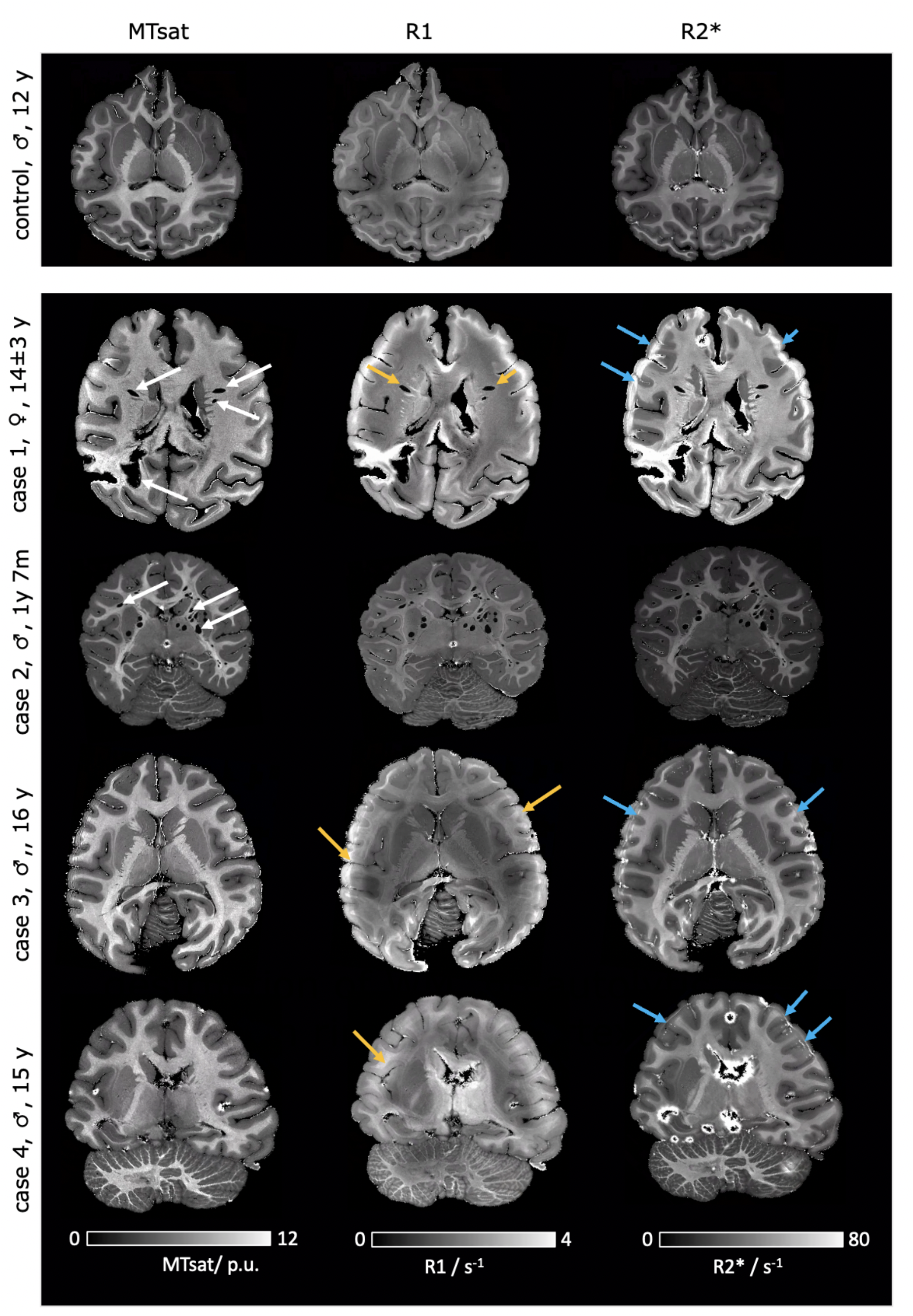
Quantitative MRI of chimpanzee brains from four confirmed cases of *Bcbva* infection (four lower rows) compared to a brain of a control individual (upper row). Three quantitative MRI parameters [magnetisation transfer saturation (MTsat), longitudinal relaxation rate (R1) and effective transverse relaxation rate (R2*)], presented in three columns, are sensitive to the tissue microstructure such as macromolecular and iron content. The signs of heterolytic damage in deep white matter (white arrows) and on the brain surface (yellow arrows) are visible in MTsat and R1 maps. The heterolytic lesions were most prominent in the cases with a higher intraparenchymal bacterial load (case 1 and case 2, see following sections and Table 1), forming large liquid-filled cavities. R2* maps show a hyperintense rim near the brain surface (blue arrows), indicating iron leaked into the tissue, putatively due to meningeal haemorrhage and superficial cortical siderosis, confirmed by Perls’s stain (Fig. 4).

In three cases (cases 1, 3 and 4), ultra-high resolution R2* maps showed a hyperintense double-banded rim in the cortex running parallel to the brain surface (Fig. 3), found to be most severe in case 3, with less intensity in cases 1 and 4. The enhanced values of iron-sensitive qMRI parameter R2* suggest elevated iron in the double-banded structure as a potential contrast source. The histological Perls’s stain for iron confirmed this suggestion, showing a double band of elevated iron content corresponding in shape and location to the hyperintensity in R2* images (Fig. 4). The iron appears finely dispersed and is not associated with specific histological structures or cell types. The bands were not visible in the unstained sections and affected only the outer parts of the brains. Their spatial distribution rather resembled the front of iron-rich compounds penetrating into the tissue from the outer surface. These abnormal iron levels and atypical distributions on the brain surface may point to superficial bleedings releasing labile iron, which then penetrated into the brain. Indeed, similar double-banded structures in the R2* maps are commonly interpreted as a sign of meningeal bleeding inducing cortical siderosis in in vivo MRI^56–60^.

**Fig. 4.**
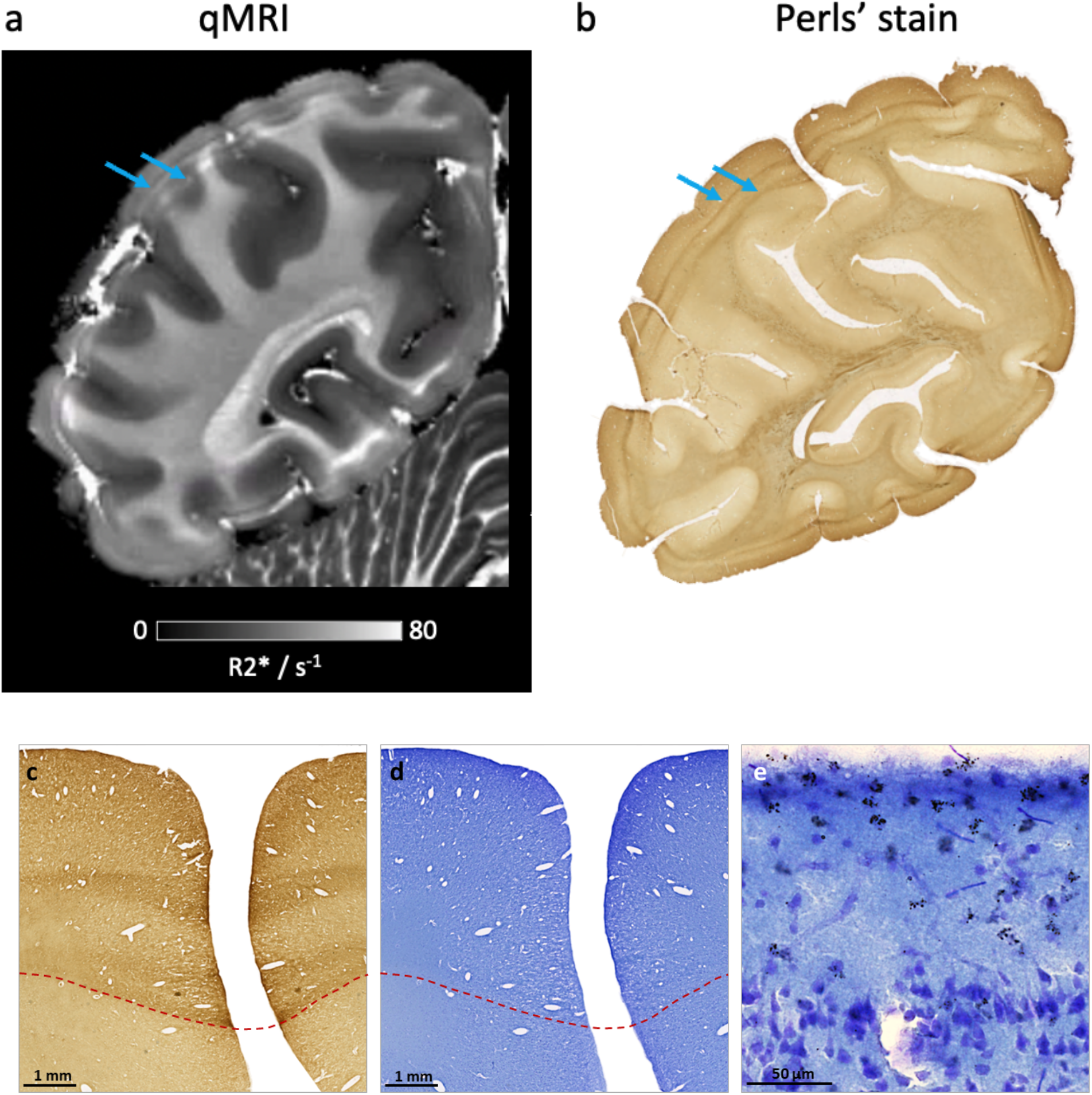
Signs of cortical siderosis caused by *Bcbva* infection in a representative case 3 revealed by quantitative R2* map (a), and histology using DAB-enhanced Perls’s stain for iron (b). R2* maps show a hyperintense double-banded rim running parallel to the brain surface, indicating iron leaked into the tissue, possibly due to meningeal bleedings causing cortical siderosis, confirmed by Perls’s stain. The characteristic double band sign in the Perls’s stain corresponds with an altered tissue structure in the Nissl stain (c,d). In high magnification the superficial cortical siderosis with dark granules in cortical layer I and II are clearly visible (e).

Cortical superficial siderosis was histologically detected in cortical layers I and II (Fig. 4e). Small dark granular deposits in layers I and II, distributed not only in the outer areas, but also in the deep sulci were found in cases 1, 3 and 4, but not in case 2 and not in the control brain (Supplemental Figure 1). The pigmented granules are already visible in the unstained sections. The presence of the granules points to *Bcbva*-induced meningeal haemorrhages consistent with the gross pathology and analogous to *B. anthracis*’ haemorrhagic meningitis in humans. Similar findings were reported in human cases with cerebral amyloid angiopathy^59^. Note that bacterial load and inflammation status of meninges could not be directly accessed, since all meninges were removed prior to start of the investigation due to requirements of a different project.

### *Bcbva* penetrates into the brain parenchyma

*Bcbva* was effectively visualised and studied in brain tissue sections using the standard Nissl stain (Supplementary Figure 2). For validation, we compared consecutive sections from the severely affected case 1, dyed using (i) Nissl stain (Supplementary Fig 2a), (ii) haematoxylin and eosin (H/E) stain (Supplementary Figure 2b), a commonly used technique in histopathology, and (iii) immunohistochemistry (Supplementary Figure 2c) employing a rabbit anti-Bacillus anthracis protein (whole cell protein) antibody. The bacteria were clearly visible with all three methods, as approximately 6 µm long and 1 µm thick elongated chains that were primarily straight and occasionally twisted. The Nissl stain exhibited superior contrast compared to H/E, showed high correspondence to immunohistochemistry and was therefore used for the investigation of all remaining cases. Nissl staining has the additional advantage of providing detailed information on cortical cytoarchitecture and separation between white and grey matter.

In all examined brain sections from the four cases, *Bcbva* bacteria were observed in both intra- and extravascular spaces. The bacterial load varied significantly between cases and brain regions (see Table 1, Fig. 5, Supplementary Fig. 3). The qualitative estimates of bacterial load performed between cases and areas showed high correspondence to the quantitative estimates of bacterial density in occipital cortex by AI data analysis (see Table 1), demonstrating the validity of the qualitative analysis. Among the cases one brain exhibited a severe (case 1) bacterial load, one brain showed a moderate level (case 2) and two brains displayed a mild overall bacterial load (case 3 and case 4).

**Fig. 5.**
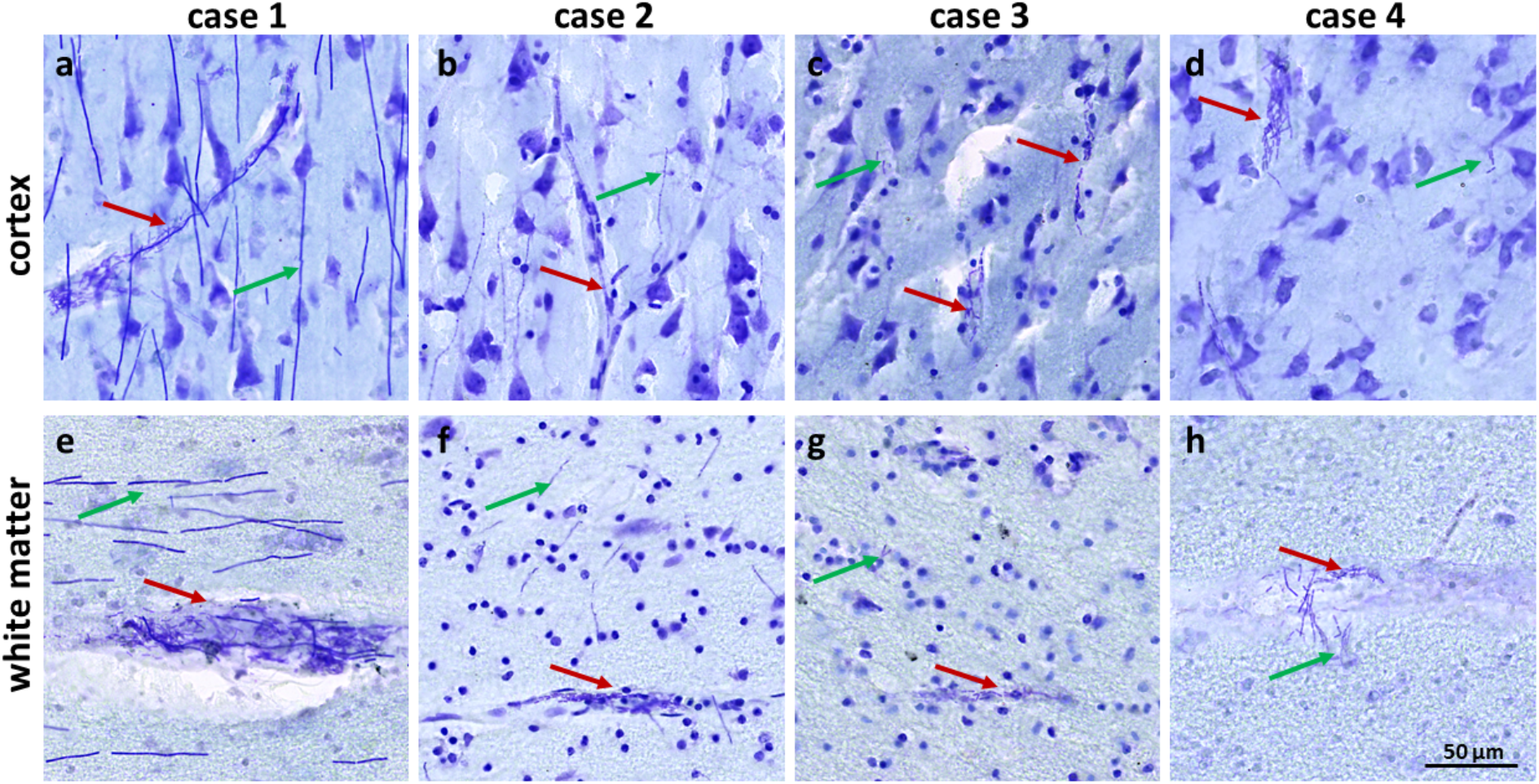
Intravascular and parenchymal distribution and density of *Bcbva*. In all cases bacteria were found intravascularly (red arrows) with moderate to high density. Invasion into the brain parenchyma could be confirmed in all cases (green arrows), however, with different densities.a Severe bacterial load was found in case 1, a moderate load in case 2, while cases 3 and 4 were mildly affected.

In the mildly affected brains (case 3 and 4), *Bcbva* bacteria were primarily found within the blood vessels in short chains, with only a few extravascular bacteria present in the cortex and white matter regions (Fig. 5 c, d, g, and h). In the brains with moderate (case 2) and severe (case 1) levels of infection, *Bcbva* bacteria infiltrated extensively into the brain tissue, forming long chains that appeared to align with the fibre orientation in the white matter and exhibited variable patterns in the grey matter. In the severely affected case 1, the bacteria had invaded the entire brain with high density. The distribution of bacteria across cortical and subcortical regions is displayed in Fig. 5.

The highest extravascular bacterial densities were found in the frontal and temporal regions and in the cerebellum, partially also in the brainstem. Striking was the highly parallel orientation of the chains in the white matter and deep cortical layer V, which seemed to follow the orientation of the myelinated fibres with chain lengths up to 200 µm. In the cortical layers I - III, *Bcbva* formed shorter chains with variable, not clearly organised patterns.

### *Bcbva* invades the brain only shortly before host death

To investigate the timeline of bacterial brain invasion, we evaluated the distribution and the morphology of micro- and astroglia as indicators of inflammation and compared them with *Bcbva* distribution patterns. Almost everywhere astrocytes exhibited predominantly normal morphology and distribution, even in regions with high *Bcbva* load, indicating none or only a low activation status (Fig. 6i, Supplementary Fig. 4). However, the GFAP immunoreactivity was increased in cortical layers I and II and around blood vessels, resulting in a strong appearance of the glia limitans and the vessel-associated astrocytes (Fig. 6g,h, Supplementary Fig. 4), indicating an early sign of astroglial activation. Microglia cells were predominantly found in a resting state, unobtrusively distributed, exhibiting ramified, non-amoeboid morphologies and only rarely appeared slightly activated (Fig. 6f, Supplementary Fig. 4). Morphologically addressable gliosis takes three or more days to develop and its absence suggests a rapid host death within days after *Bcbva* infiltration of the central nervous system^61,62^.

**Fig.6.**
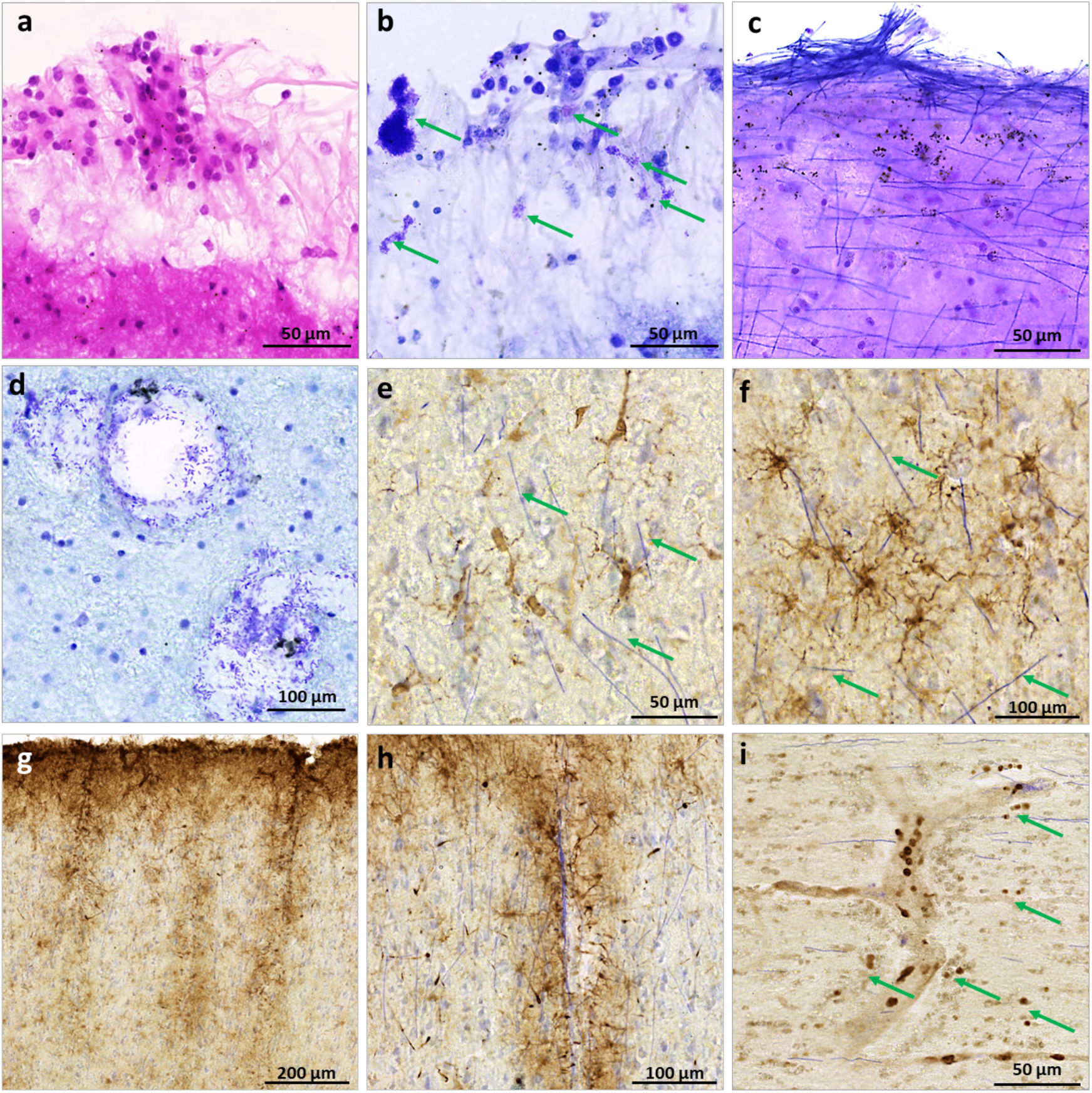
Histological evidence of *Bcbva*’s invasion of and effects on the central nervous system (case 1 shown): (a,b) Pia mater adjacent to the frontal cortical surface with leukocytes, indicating meningitis, visualised by H/E stain in (a) and aggregated bacilli (arrows) visualised by Nissl stain (b); (c) Superficial cortical siderosis in the frontal cortex layer I-II with haemosiderin particles and dense conglomerates of bacilli on the cerebral surcface (H/E stain), (d) Blood vessels with Bcbva infiltration into the parenchyma through a broken blood brain barrier (Nissl), (e) No morphological signs of microglia activation was detected (rabbit anti-Iba1 antibody, DAB, Nissl), (f) Also, astrocytes were regularly distributed and appeared morphologically unaltered (rabbit anti-GFAP, DAB, Nissl), (g,h) However, enhanced glial fibrillary acidic protein (GFAP) immunoreactivity was apparent in the glia limitans in the outer cortical layers and close to walls of penetrating blood vessels, revealing early stages of astroglial activation (rabbit anti-Iba1 antibody, DAB, Nissl) (i); MPO+ leucocytes were mostly detected inside vessels and rarely entered the parenchyma (arrows) (rabbit anti-MPO antibody, Nissl).

Myeloperoxidase (MPO) positive cells (neutrophilic granulocytes, monocytes) were predominantly found in blood vessels, rarely invading the brain tissue (Fig. 6i, Supplementary Fig. 4), providing further support for a rapid host death.

### *Bcbva* breaches blood-brain barriers

To identify the path of bacterial brain invasion we studied the integrity of brain-blood and blood-CSF barriers, by looking at bacterial distribution on the mesoscale and the presence of blood macrophages in the parenchyma.

In many regions *Bcbva* chains were piled up on the brain surface and abundant in the upper cortical layers and adjacent pia mater, partially in dense bacterial plaques (Fig. 6b,c). This indicates a primary disintegration of the arachnoidal barrier in the bacteraemic host with secondary parenchymal invasion from the subarachnoid space. Additionally, however to a much lesser extent, bacteria were visualised in the Virchow-Robin spaces and transgressing into the adjacent brain tissue, evidencing a disintegration of the blood-brain barrier’s neurovascular unit (Fig. 6d).

### *Bcbva* degrades neuronal extracellular matrix

To assess *Bcbva’s* impact on parenchymal tissue integrity, perineuronal nets (PNNs) were stained via anti-aggrecan immunohistochemistry. PNNs are a specialised form of cortical ECM consisting of a mesh-like arrangement of proteins and sugar molecules ensheathing certain neurons, with aggrecan constituting a key component of healthy neurons^52^. A clear sign of progressive tissue degradation in the *Bcbva* infected brains is evidenced by the loss of aggrecan positive PNNs (Fig. 7). The severe reduction of aggrecan positive PNNs between the cases exemplarily shown in the occipital cortex was strongest in cases with the higher load of *Bcbva* (Table 1).

**Fig. 7.**
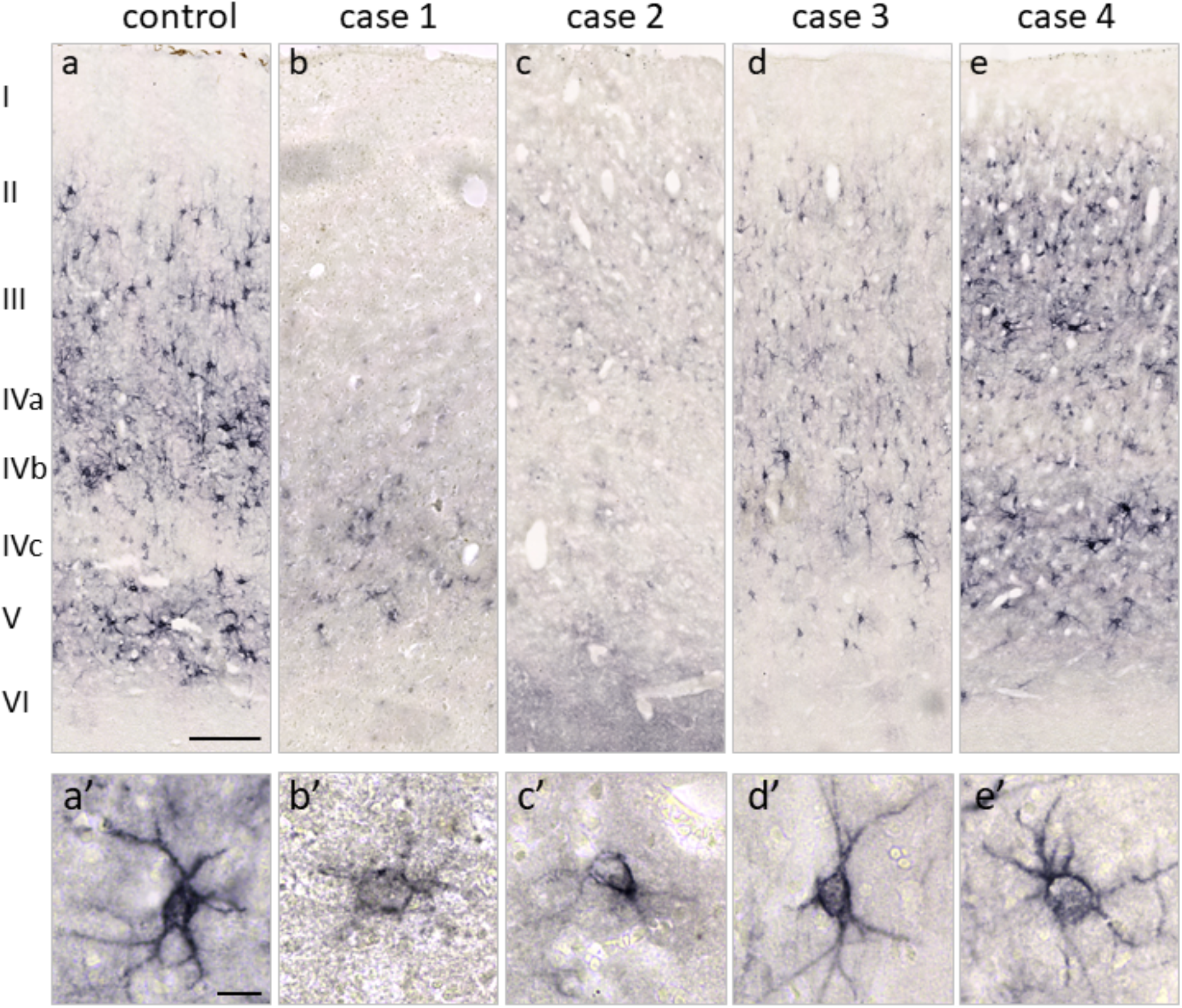
Laminar distribution of the major extracellular matrix (ECM) protein aggrecan (acan) in the occipital lobe (chimpanzee analogue of Brodmann Area 17, primary visual cortex V1) of *Bcbva* infected cases and one control. (a) Acan as an established marker for perineuronal nets indicated many perineuronal nets (PNNs) in layer III, IVa and V. Fewer PNNs were seen in layer VI, whereas layer I, II and IVc are virtually devoid of acan positive PNNs. (a, a’) This pattern reveals the expected distribution of PNNs in a healthy hominoid brain with regularly structured PNNs. (b,c,d,e) A severe reduction of PNNs is visible in severely affected cases 1 and 2. Less reduction was observed for Cases 3 and 4 with lower bacteria load. Scale bars (a)-(e): 200µm, (a’)-(e’): 20µm.

The number of aggrecan positive PNNs was reduced by up to 96 % in the most severely *Bcbva* infected case 1 compared to the non-infected control brain (Table 1).

## Discussion

### General neuroinvasiveness, genetic homogeneity

We demonstrated that *Bcbva* is neuroinvasive in one of our closest living relatives, chimpanzees, and hence likely to have similar properties in humans. These insights were only facilitated by a multidisciplinary and multimodal approach and the unique and ethically sustainable sampling of high quality brain specimens under challenging field conditions in the Taï NP, requiring continuous behavioural observation of habituated chimpanzee groups and the presence of on-site veterinarians^31^.

We observed direct and indirect evidence for meningitis, analogous to *B. anthracis*-induced anthrax in humans^18^. Furthermore, our findings demonstrate that *Bcbva* penetrated the brain parenchyma shortly before death, resulting in the degradation of brain tissue and digestion of neuronal extracellular matrix (ECM) in all four cases. This is in contrast to *B. anthracis*, which, despite causing severe meningitis, rarely invades the deeper brain tissue of either humans and NHP^19,63^. Thus, based on our results, we can conclude that *Bcbva* may be more neuroinvasive in terms of parenchymal invasion, compared to its more extensively studied cousin.

Genetical analysis for two out of four studied cases did not reveal any virulence-altering mutations, indicating that their ability to cause CNS disease is representative for *Bcbva* (Fig. 2).

### Blood-brain-barrier breaches resulting in meningeal and parenchymal bacterial spread

We showed that *Bcbva* is able to breach the blood-barriers, which normally hinder dissemination of bacteria into CNS compartments. These entail the blood-brain barrier formed by the neurovascular unit (NVU), preventing pathogen dissemination into the brain tissue at the parenchymal vessel interface^64^ and the blood-cerebrospinal fluid barriers at the meningeal vessels (arachnoid barrier) as well as at the choroid plexus^65^.

Studies have shown that classical *B. anthracis* successfully penetrates the blood-cerebrospinal-fluid barrier and thrives within the cerebrospinal fluid (CSF), as evidenced by the presence of abundant bacilli distending the subarachnoid space^66,67^.

Similarly, our observations indicated the presence of *Bcbva*-induced meningitis in all four investigated cases. In case 1, direct histological examination revealed the presence of bacilli and leukocytes within pia mater residues (Figure 6a, b, c). This microscopic evidence was supported by macroscopic observations of congested pial vessels and meningeal haemorrhages (Fig. 1). Additionally, cortical siderosis, detected using qMRI (Fig. 3 and Fig. 4), suggested the occurrence of meningeal haemorrhage (Charidimou et al. 2020; Shams et al. 2016; van Veluw et al. 2020). We note that direct investigation of pia mater was only possible on fragments of pia mater, since it was removed before MRI scans for other projects.

Analogous to *B. anthracis*, *Bcbva* likely overcomes the cerebrospinal fluid barrier with the help of anthrax toxin mediated vascular damage^68,69^, as well as the proteolytic activity of InhA1, cleaving zonula occludens-1 proteins and thus degrading tight junctions^70^. Additionally, the virulence factor AtxA has been shown to play a crucial role in CNS invasion^71^.

Despite massive meningeal involvement, *B. anthracis* bacilli are rarely visualised within the adjacent brain parenchyma^19^ and if so, they are predominantly found in perivascular spaces rather than in the neuropil^63^. Consequently, some authors concluded that *B. anthracis* causes meningitis rather than encephalitis^19,72^, which is similar to other extracellular neuroinvasive bacteria like *Neisseria meningitidis*, *Streptococcus pneumoniae* and *Haemophilus influenzae* ^73^.

In contrast, *Bcbva* was observed within the brain parenchyma in all four cases (Table 1, Fig. 5), indicating a greater ability to transgress the glia limitans, which is formed by astrocyte endfeet plus their associated basal lamina and constitutes as a secondary barrier to bacterial invasion of the neuropil^74^. The factors that enable *Bcbva* to invade the brain parenchyma so effectively require further investigation. One potential contributing trait is *Bcbva’*s flagellar motility, as opposed to the non-motile *B. anthracis*^11^.

The observation of highest bacterial density in layer 1 of the cerebral cortex suggests parenchymal infiltration predominantly from the subarachnoid space. But additional invasion via parenchymal vessels was detected (Fig. 6h, i).

### Extracellular matrix degradation

Upon invasion, *Bcbva* extensively degrades CNS-tissue as evidenced macroscopically by the lytic cavities visible in quantitative MRI maps (Fig. 3) and confirmed by histology, as well as on a finer structural level by the digestion of perineuronal nets (PNNs) (Fig. 7). PNNs form a mesh-like structure around neurons composed of ECM components like hyaluronan, chondroitin sulfate proteoglycans and link proteins^75^ and are likely cleaved by *Bcbva’s* protease secretion. The secretome of *Bcbva* and *B. anthracis* were shown to be largely congruent^76^, and for the latter a major portion of secreted proteins are proteases, foremost the two-neutral zinc metalloproteases InhA and Npr599^77^. InhA exhibits a low substrate specificity, cleaving a variety of extracellular matrix (ECM) proteins and acts as multifactorial pathogenic factor aiding with tissue degradation, breaking barriers and interfering with immune functions^78^.

### Propagation in Tissue

This degradation of the extracellular matrix plausibly shapes *Bcbva’s* within tissue spread, resulting in parallel, periaxonal bacilli propagation patterns within the white matter, and a more dispersed growth pattern in the cerebral cortex (Supplementary Fig. 3). Additionally to intraparenchymal multiplication in straight chains, also twisted cell morphologies have been observed. This growth form has been described in-vitro on culture media^79^ and its occurrence in vivo may help in etiological diagnostic assessment.

### Timeline of Invasion

The absence of an obvious structural gliosis is remarkable and provides valuable insights into the timeline of bacterial brain invasion. Normally, when the CNS is subjected to an inflammatory stimulus, both astrocytes and microglia are activated, characterised by cellular restructuring. Astrocytes become hypertrophic and their processes extend, while microglia undergo a transition from a resting (ramified) morphology to an amoeboid one^80,81^. The observed multifocal cortical and perivascular increase in GFAP levels suggests an early stage astrocyte activation, since GFAP gene expression can increase as early as 1 hour after an inflammatory stimulus^82^. However, cellular restructuring takes longer to manifest. In a mouse model of traumatic brain injury, for example, hypertrophic astrocytes were only visible 3 days after the initial inflammatory insult^62^. The same applies to microglia responses which usually require hours to days to fully develop^61^. Therefore, the lack of structural gliosis argues for a rapid death of the chimpanzees following CNS invasion, likely within less than 48 hours.

This timeline is supported by little intraparenchymal leukocyte recruitment^83^, although in the residual of the meninges and the vessels leucocytes were visualised (Fig. 6 a,b,i). Combined with the observation of greater intravasal than extravasal bacterial load (except for case 1), this suggests initial bloodborne subarachnoid pathogen spread with secondary parenchymal propagation across the glia limitans. *Bcbva* multiplies rapidly, comparable to *B. anthracis*, which features doubling times of 30-60 minutes^84^, reached at a late infection stage once the host immune system has been overwhelmed^28,85^. This explains the excessive pathogen load (above all in case 1) despite a per-acute course of disease.

The variation in bacterial density observed in different cases might be attributed to various factors. These include differences in the individual host’s immune status and susceptibility to *Bcbva*, as well as the initial infection dose and/or the extent of bacterial propagation after death and before fixation^86^.

### Clinical symptoms

The observed sudden chimpanzee deaths are comparable to human *B. anthracis* cases with CNS involvement, in which 75% of the patients die within 24 hours after initial presentation^20^. It is challenging to determine the exact onset of clinical signs from behavioural observations in chimpanzees as wild animals typically conceal weakness as long as possible^87^. Although not observed for the presented cases, other *Bcbva*-infected chimpanzees have been reported dying in a ventral position holding their heads, suggesting severe headache (unpublished observations from TCP health monitoring programme by FL). This is consistent with *B. anthracis* involving the CNS^22^. Behavioural observations prior to death were available in two investigated cases (2 and 4), with absence of clinical signs in the young individual (2) 1.5 hours prior to its sudden death and progressively altered behaviour for case 4 with sleepiness, isolation from the group and increased resting periods in the preceding two days. These clinical signs might be partially accounted for by involvement of the CNS but are by no means specific.

### Implications for human spillover and treatment

Our findings indicate that a human *Bcbva* infection would likely lead to meningitis and encephalitis after bacteremic spread with a very poor prognosis. Nevertheless, the precise rate at which infections progress to a systemic and septicemic state remains uncertain. One study revealed a 10 % seropositivity for *Bcbva* antibodies among people residing near Taï NP, suggesting frequent non-lethal infections. In contrast, a separate study conducted on wildlife found limited seropositivity, implying a high lethality associated with *Bcbva* infections given its significance as a driver of wildlife mortality. A human *Bcbva* case involving the CNS would entail management analogous to a *B. anthracis* meningitis case including intravenous fluoroquinolone administration and one or more agents with good CNS penetration (e.g. rifampicin)^21^. As opposed to *B. anthracis*, penicillin is not advised as central African *Bcbva* isolates may be resistant^11^. Early initiation of treatment is critical as produced toxins sustain detrimental effects even after bacteria have been cleared, especially since antibody-based therapeutics (e.g. raxibacumab) cannot neutralise already internalised toxins^88^. Etiologic disease diagnosis and especially early diagnosis are challenging in rural African regions, since general health care infrastructure is usually poor^89,90^. Therefore, especially in regions with bushmeat consumption, local public health sensitisation and diagnostic training pose the first critical steps towards the mitigation of potential human zoonotic *Bcbva* mortality. Further it highlights the importance of effective disease surveillance in wildlife to detect heightened risk of spillover early-on in the context of a One Health approach.

### Limitations

One of the study’s major strengths, the opportunistic, ethical sampling approach, intrinsically comes with limitations such as the lack of standardised infection doses and *postmortem* intervals. *Bcbva* is facultatively anaerobic and may multiply even after animal death. This might have influenced the bacterial density and tissue integrity. Still, the core statement regarding pronounced *Bcbva* neuroinvasiveness remains unaffected. Indeed, the initialised inflammatory GFAP response indicated *premortem* penetration into the parenchyma. The absence of secondary, opportunistic bacteria in all cases showed that *postmortem* time before fixation was not sufficient for massive *postmortem* bacterial invasion. The fact that the blood-brain barrier remains intact for a certain period even after death^91^ makes *postmortem* bacterial propagation and growth in brain tissue unlikely. Another limitation is the lack of meningeal tissue, resulting in a lack of detailed histological assessment of meningeal involvement. Therefore we relied on gross pathology and indirect signs of siderosis to infer the presence of meningitis. Further studies, including histological analysis of the larger meninges samples, are needed to confirm our findings.

## Acknowledgements

We are grateful for research and brain extraction permissions in Sierra Leone to Environmental Protection Agency; in Côte d’Ivoire to the Ministère de l’Enseignement Supérieur et de la Recherche Scientifique and the Ministère de Eaux et Fôrets in Côte d’Ivoire. We are grateful to the Centre Suisse de Recherches Scientifiques en Côte d’Ivoire for their logistical support, the staff members of the Taï Chimpanzee Project and the Tacugama Chimpanzee Sanctuary for their support and assistance in collecting the samples and data. We especially thank Christina Kompo for adeptly handling import and export permits and shipping.

## Appendix 1

EBC Consortium:

Tobias Gräßle (1,2), Ariane Düx (1,2), Carsten Jäger (3,4), Evgeniya Kirilina (3), Jenny Jaffe (2,8), Penelope Carlier (8), Andrea Pizarro (5), Anna Jauch (3), Katja Reimann (4), Angela Friederici (3), Philipp Gunz (11), Ilona Lipp (3), Bala Amarasekaran (5), Roman M. Wittig (6,7,8), Catherine Crockford (6,7,8), Nikolaus Weiskopf (3,9,10), Fabian H. Leendertz (1,2), Markus Morawski (3,4)

(1) Ecology and Emergence of Zoonotic Diseases, Helmholtz Institute for One Health, Helmholtz Centre for Infection Research, Greifswald, Germany; (2) Epidemiology of Highly Pathogenic Microorganisms, Robert Koch Institute, Berlin, Germany; (3) Department of Neurophysics, Max Planck Institute for Human Cognitive and Brain Sciences, Leipzig, Germany; (4) Paul Flechsig Institute - Center of Neuropathology and Brain research, Medical Faculty, University of Leipzig, Germany; (5) Tacugama Chimpanzee Sanctuary, Freetown, Sierra Leone; (6) Evolution of Brain Connectivity Project, Max Planck Institute for Evolutionary Anthropology, Leipzig, Germany; (7) Ape Social Mind Lab, Institute for Cognitive Sciences Marc Jeannerod, UMR CNRS 5229, University Claude Bernard Lyon 1, Bron, France; (8) Taï Chimpanzee Project, CSRS, Abidjan, Côte d’Ivoire; (9) Felix Bloch Institute for Solid State Physics, Faculty of Physics and Earth Sciences, Leipzig University, Leipzig, Germany ; (10) Wellcome Centre for Human Neuroimaging, UCL Institute of Neurology, University College London, London, UK; (11) Max Planck Institute for Evolutionary Anthropology, Leipzig, Germany.

## Supplementary Information

### Ultra-high resolution multi-parametric MRI acquisition

Quantitative maps of MRI parameters were obtained using a multi-parametric mapping method^1–3^. A multi-echo 3D gradient echo sequence with an isotropic resolution of 300 μm (matrix: 432 x 378 x 288; readout bandwidth of 331 Hz/pixel; repetition time TR = 70 ms; 12 equidistant echoes with echo time TE1-12=3.63ms-41.7 ms with a bipolar gradient readout, △TE = 3.56 ms) was acquired using three different sets of parameters for proton-density-weighted (PD_W_), magnetisation-transfer-weighted (MT_W_) and longitudinal relaxation time-weighted (T1_W_) images. PDW- and MTW-images were acquired using excitation flip angles (FA) of 18°, the T1_W_ image was acquired with FA=84°. Gaussian MT saturation pulse was applied at 3 kHz offset-off resonance frequency and FA=700°. Radio frequency (RF) excitation transmit (B1^+^) field maps were acquired using a simultaneous spin and simulated echo measurement^1,3^ with an isotropic resolution of 4 mm, with parallel imaging acceleration factor 2 in both phase encoding directions. Maps of the static magnetic field (B_0_) were acquired with the isotropic resolution of 2 mm and following acquisition parameters: TR = 1020 ms, TE = 10 and 11.02 ms, FA = 20°. To correct RF receive bias (B1^-^) field low resolution (2.1 mm) T1_W_ images were acquired with the 32-channel receive coil vs the image acquired with the RF transmit element used also for receive.

---

**Supplementary Table 1:** Case descriptions of four chimpanzees with confirmed infection of Bcvba and one control case. PMI - *post mortem* interval before fixation. F=female, M=Male. *Age estimate.

| Case Nr. | Sex | Age | PMI | Clinical signs | Regions investigated with histology |
| --- | --- | --- | --- | --- | --- |
| 1 | F | 14±3 y* | 24 h | Unknown | 9 coronal blocks right<br>4 brainstem blocks<br>cerebellum |
| 2 | M | 1.7 y | 24 h | Seen with no symptoms 1 hour before death | frontal left/right<br>central left/right<br>occipital left/right<br>brainstem |
| 3 | M | 16 y | < 16 h | Normal general condition 5 days before death | frontal right<br>central right<br>occipital right<br>brainstem |
| 4 | M | 15 y | < 12 h | Lethargy, isolation from the group 3 days before death | occipital right |
| Control | M | 12 y | 6h | None | frontal left/right<br>central left/right<br>occipital left/right |

**Supplementary Table 2:**
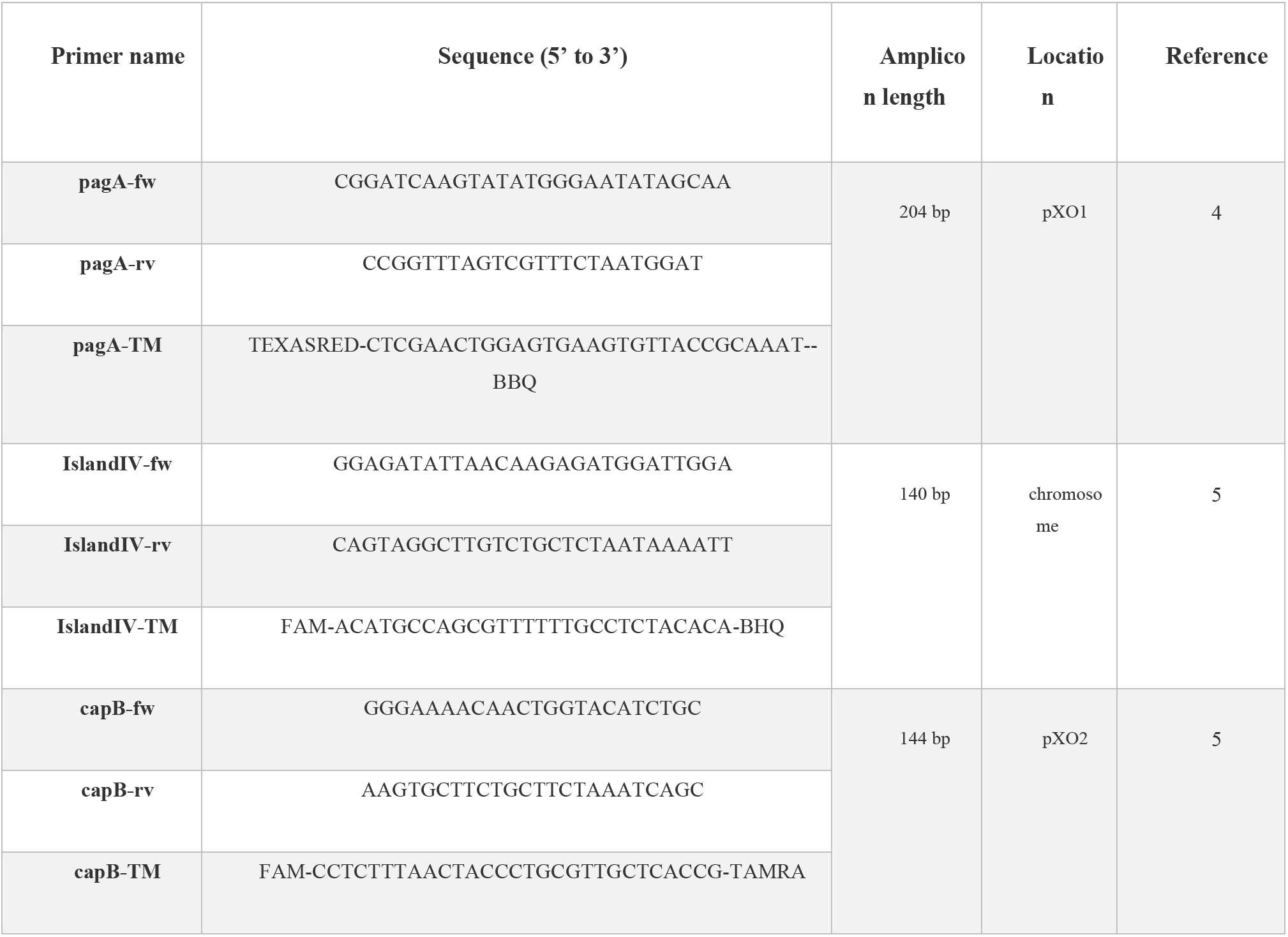
qPCR primers for *Bcbva* detection.

https://docs.google.com/spreadsheets/d/1IcHgEdFVIq2rQEWbIoxqadgWu9pDWkahJlYPdZSsFzw/edit?usp=sharing

**Supplementary Table 3:**
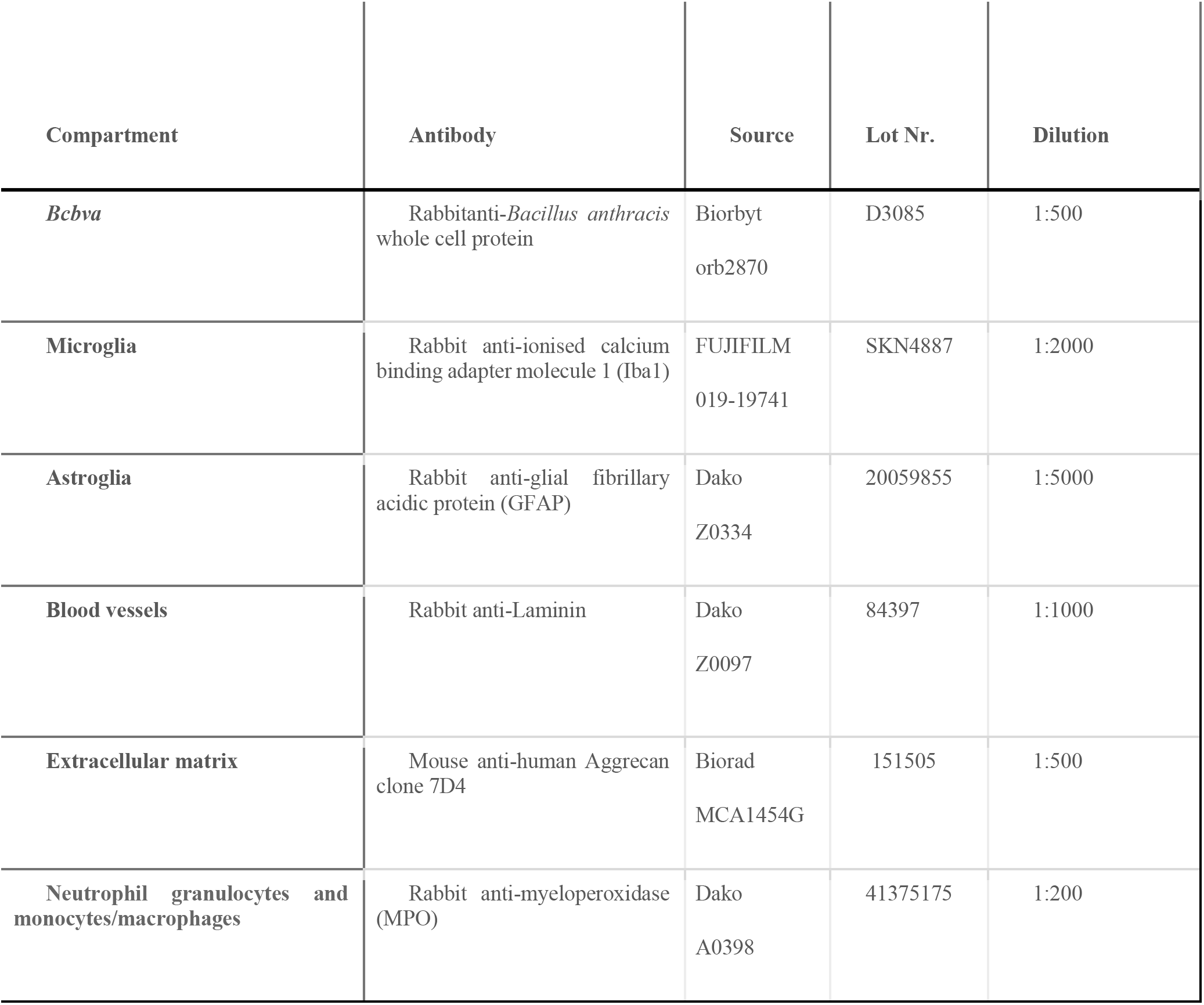
Primary antibodies used for staining of specific compartments.

**Supplementary Fig. 1:**
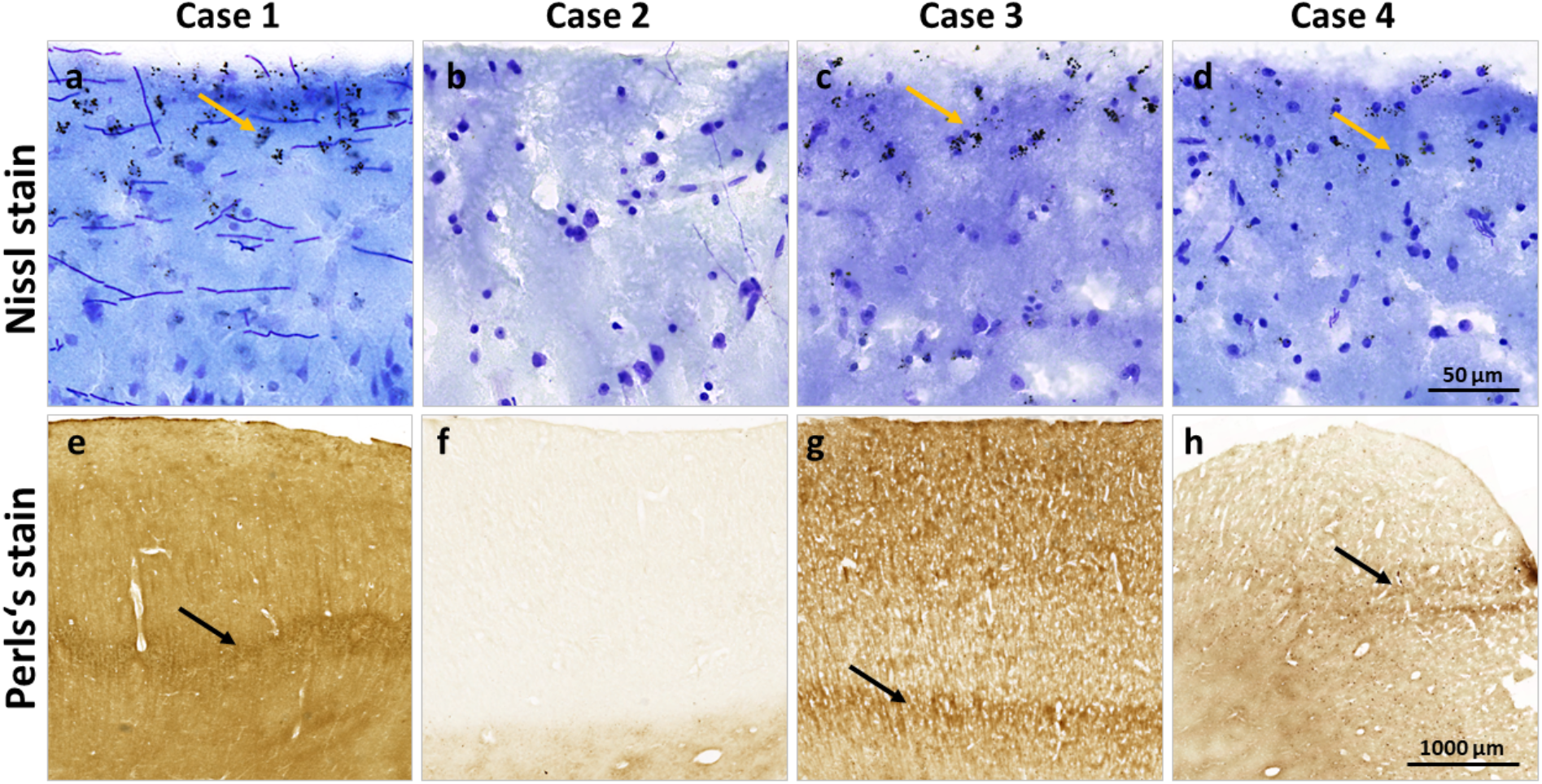
Superficial cortical siderosis and iron-rich double-banded rim was observed in three out of four cases. Superficial cortical siderosis with dark granules in cortical layers I and II is present in cases 1, 3 and 4 (a, c, d), but not in most regions of case 2 (b). The iron-rich double-banded rim visible in R2* maps also appears in the Perls’s stain in cases 1, 3 and 4 (e, g, h). Case 2 reveals an overall low iron level due to the young age of the specimen and does not show the rim in both R2* maps and Perls’s stain (f).

**Supplementary Figure 2:**
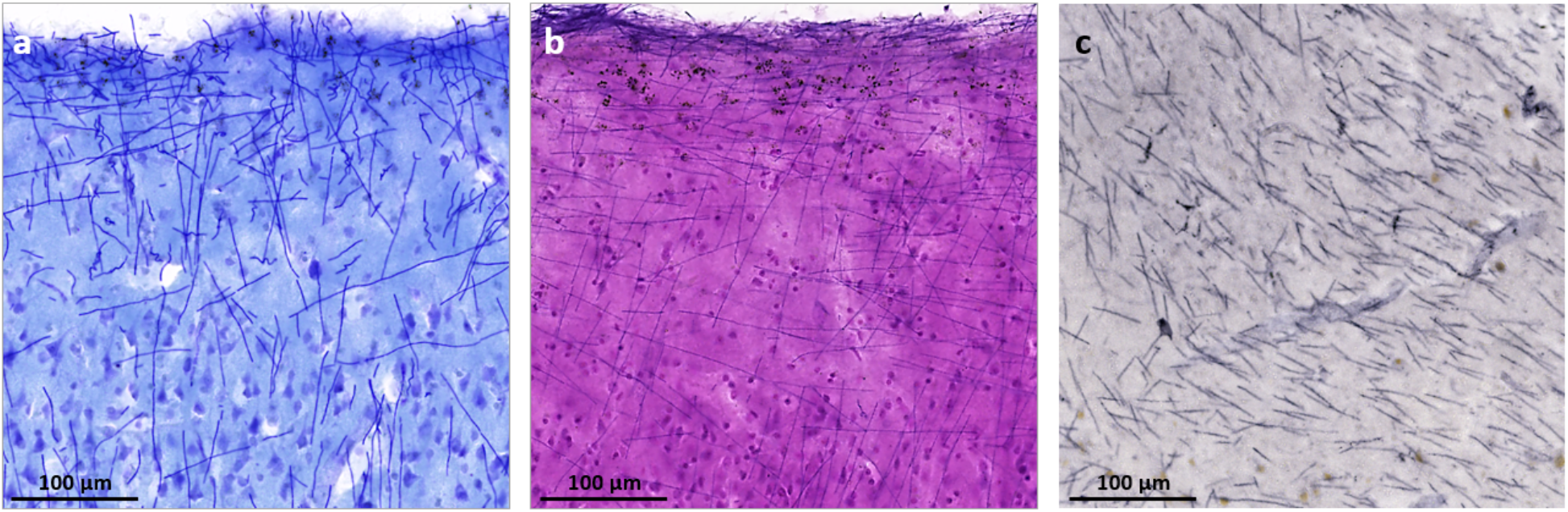
Visualisation of *Bcbva* in case 1 with (a) Nissl (cresyl violet) stain, (b) haematoxylin/eosin stain and (c) validation by immunohistochemistry with rabbit anti-*B. anthracis* protein (whole cell protein). Characteristic elongated chains of the pathogen were visible in H/E stain and Nissl stain and corresponded well in length, thickness and distribution to the gold standard immunohistochemical stain.

**Supplementary Figure 3:**
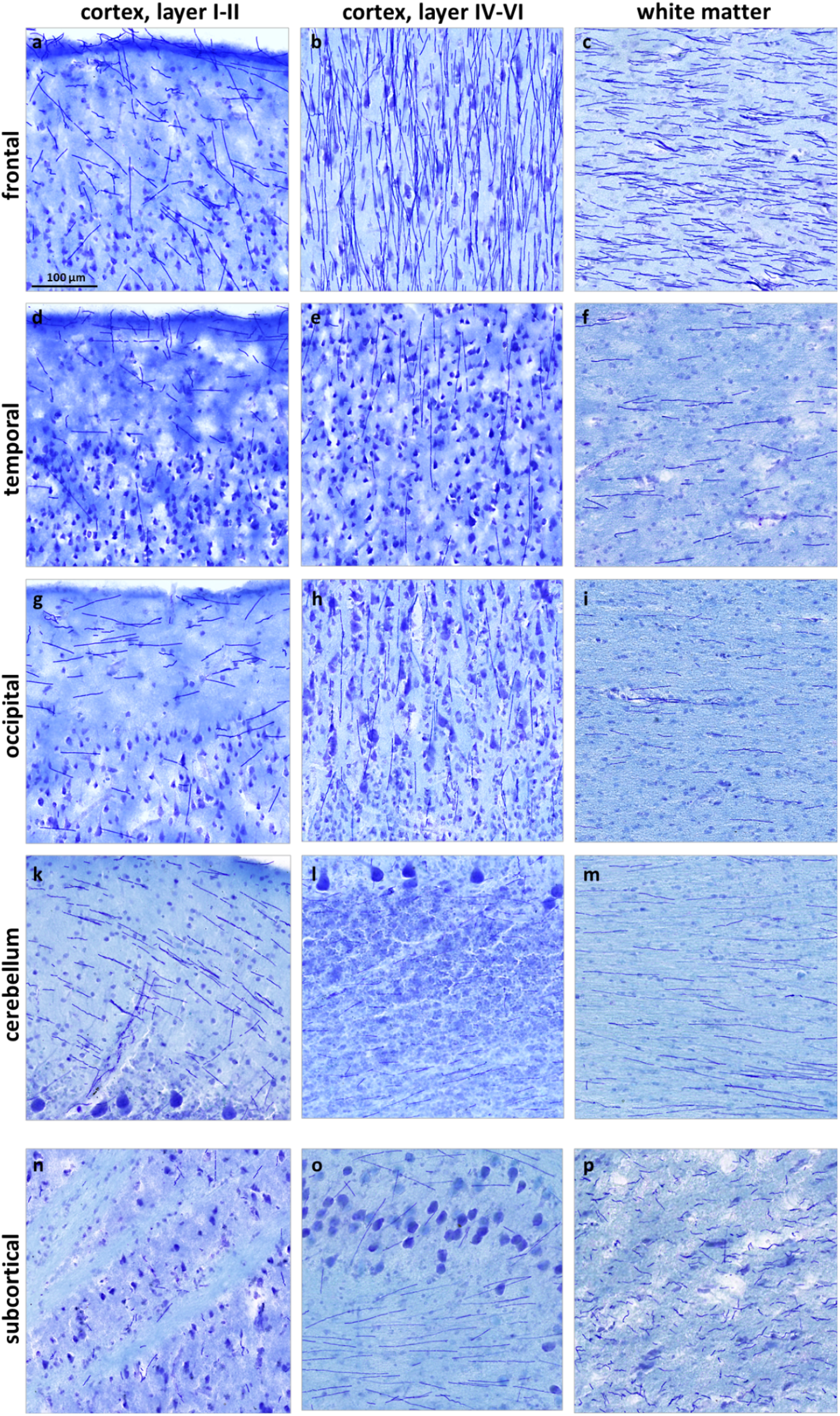
Distribution patterns and density of *Bcbva* in the most severely affected case 1. *Bcbva* chains were detected across the entire brain: (a) frontal cortex, layer I-III (b) frontal cortex, layer V (c) frontal lobe, white matter (d) temporal cortex, layer I-III (e) temporal cortex, layer V (f) temporal lobe, white matter (g) occipital lobe, V1, layer I-III (h) occipital lobe, V1, layer IVa, IVb (i) occipital lobe, white matter (k) cerebellum, molecular layer (l) cerebellum, granular layer (m) cerebellum, arbor vitae (n) striatum (o) brainstem, inferior olivary nucleus (p) brainstem, pons.

**Supplementary Figure 4:**
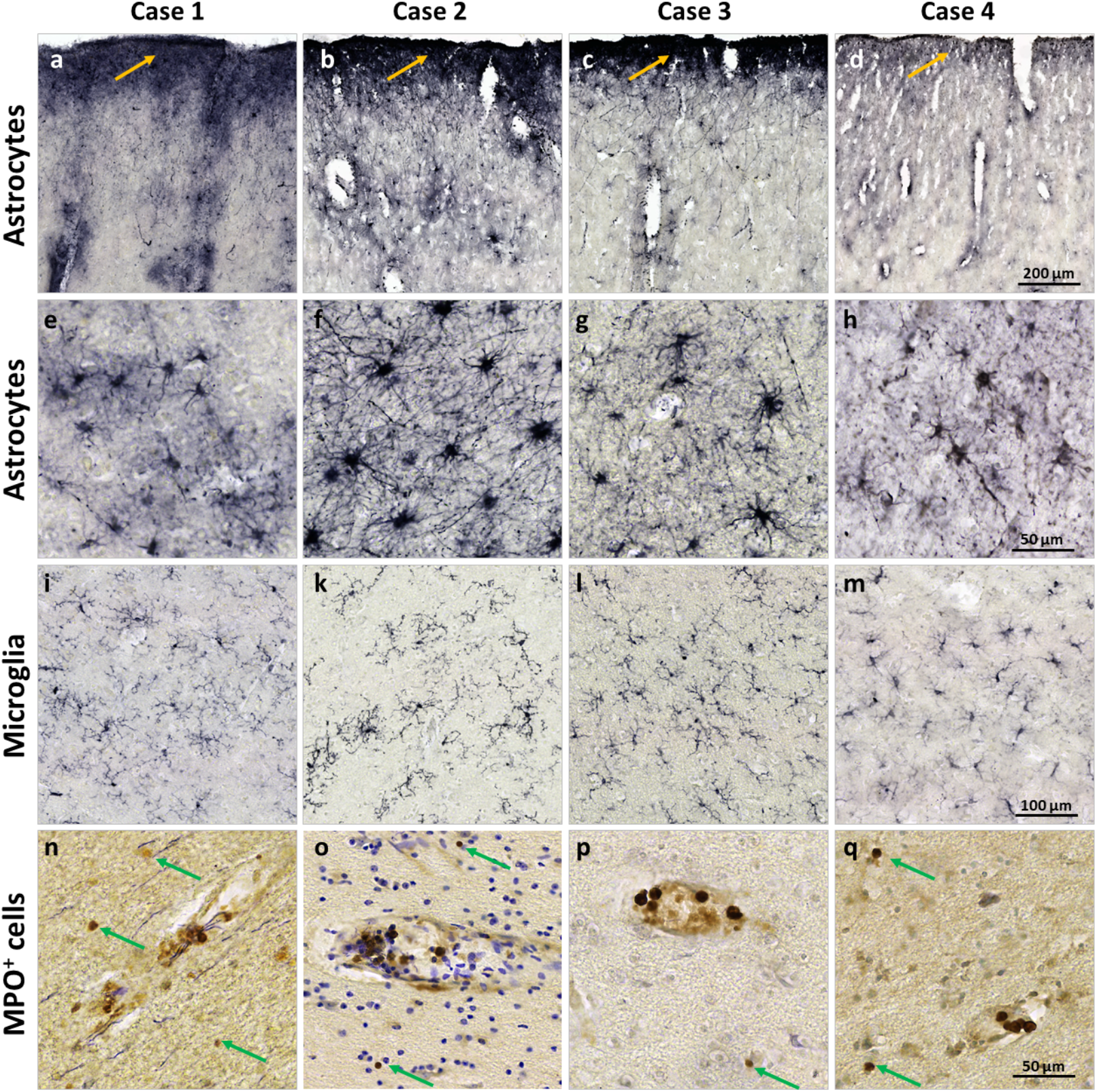
Glial reaction and MPO^+^ leukocyte invasion for all cases. Enhanced GFAP immunoreactivity is present in all cases in the glia limitans on the brain surface as well as around penetrating blood vessels (a - d; rabbit anti-GFAP, Ni-DAB). White matter astrocytes are regularly distributed and do not reveal any morphological changes, indicating no signs of activation (e - h; rabbit anti-GFAP, Ni-DAB). Microglia appears regularly shaped and distributed (i - m; rabbit anti-Iba1 antibody, DAB). MPO+ leucocytes were found in blood vessels and rarely infiltrated in the brain tissue (n - q; rabbit anti-MPO antibody, Nissl; MPO+ leukocytes outside vessels are indicated by arrows).

